# Risks inherent to mitochondrial replacement

**DOI:** 10.1101/015628

**Authors:** Edward H. Morrow, Klaus Reinhardt, Jonci N Wolff, Damian K Dowling

## Abstract

The UK Government has recently been debating whether or not to legislate to allow *mitochondrial replacement* (MR) to be used in the clinic. However, we are concerned that some of the science of MR has been misunderstood, or otherwise given only fleeting consideration. We set out our arguments below and offer a way forward to ensure that MR can safely deliver the health benefits it promises for those suffering from mitochondrial-related diseases

Recent innovations that enable mitochondrial DNA (mtDNA) mutations to be eliminated from the germline, by replacing mutated mitochondria within an oocyte with mitochondria from a healthy donor female^1–3^, offer hope for the eradication of several debilitating and lethal mitochondrial diseases. The potential for clinical application of MR has received widespread support^4,5^, but has also provoked safety and ethical concerns from the public and biomedical practitioners^4,6^. In addition to currently addressed safety concerns related to technical details of the procedures^7^, a further safety concern exists that cannot be easily addressed by methodological refinements. Embryos produced by all variants of MR (pronuclear transfer; maternal spindle transfer; polar body transfer) will acquire genetic material from three different individuals (nuclear DNA from the prospective parents, and mtDNA from a donor female), and some of these novel combinations of genetic material may not be fully compatible with one another (i.e. may be mismatched). For example, various combinations of donor mtDNA and recipient nuclear genomes have experimentally been shown to negatively affect offspring health and fitness in vertebrate and invertebrate models, even though the donated mitochondria were putatively healthy^8^. This evidence has, however, been suggested to have low relevance to humans^3,7,9,10^ for three proposed reasons (Box 1). Here we address each of those reasons, and explain why none of them refute compellingly the potential for mitochondrial-nuclear (mito-nuclear) mismatches to affect the outcomes of MR in humans.

## Box 1. Three proposed reasons why MR should not result in alterations of human phenotypes

### Reason 1. MR, like sexual reproduction, randomly shuffles mitochondrial and nuclear genomes each generation

This is based on the argument that sexual reproduction results in the random mixing of two parental genomes. Thus, under sexual reproduction, the father’s haploid genome is as evolutionarily ‘foreign’ to the mother’s mtDNA, as will the mother’s nuclear genome be to a donor’s mtDNA under MR^9^. Under the additional assumption that the mito-nuclear combinations found in the offspring are a random subset of those determined at fertilization (i.e. the absence of selection is assumed), there will be little scope for high performing mito-nuclear allelic combinations to be preserved across generations. MR has therefore been described as being equivalent to sexual reproduction, in terms of generating healthy offspring containing novel combinations of mitochondrial and nuclear alleles. In section 1, we explain why the process of co-transmission of mtDNA and maternal nuclear DNA, coupled with selection, renders this proposed reason unconvincing.

### Reason 2. Genetic diversity in humans is too low to cause incompatibilities

It has been suggested that mito-nuclear mismatches are unlikely to occur in humans because the genetic diversity within the human population is so small that any disruptions will be negligible^7^. It has been argued that mito-nuclear compatibility should be widespread, given that humans are “a freely interbreeding species”^10^. In section 2, we outline why the potential for mito-nuclear incompatibilities in humans remains a credible possibility.

### Reason 3. Incompatibilities do not occur in non-human primates

Empirical data in a primate model^3^ has been used as evidence that mito-nuclear mismatching will not occur, or will not be important, in humans. This reasoning is based on the production of four healthy male macaques born to three mothers, following MR-assisted IVF attempts on twelve mothers^3^. The individuals were apparently derived from two distinct, although unspecified^1^, sub-species of *Macaca mulatta*^9^. In section 3, we outline why the macaque studies, to date, do not provide a strong base on which to dispel concerns regarding mito-nuclear incompatibilities manifesting in humans.

## 1. MR is more likely than sexual reproduction to disrupt coevolved mito-nuclear genetic combinations

### 1.1 Co-transmission

During sexual reproduction, but not during MR, offspring invariably receive an entire haploid copy of the nuclear g enome from their mother, alongside their maternally-inherited mtDNA. In other words, mitochondrial alleles co-transmit with 50% of the autosomal nuclear alleles in 100% of the cases (and with two-thirds of the X-chromosome linked alleles, since females carry two copies of the X-chromosome and males carry only one). In contrast to sexual reproduction, MR can create entirely novel allelic combinations of mito-nuclear genotypes, because the mtDNA has been donated from a third-party (the donor female) – thus the co-transmission rate between the patient’s nuclear DNA and the donated mtDNA is 0%.

### 1.2 Selection

The co-transmission of mtDNA and nuclear alleles facilitates the preservation of high performing (coevolved) combinations, across generations. The greater the percentage of co-transmission between mtDNA and nuclear DNA, the higher the potential for mito-nuclear co-adaptation. In natural conceptions, embryos carrying better-performing mito-nuclear allelic combinations may be more likely to survive through development, to reach reproductive age, and ultimately to successfully reproduce. Because the best-performing combinations may be more likely to be passed on, coadapted mito-nuclear allelic pairings are likely to be preserved across generations within any particular maternal lineage. Similarly, germline selection against incompatible mito-nuclear combinations might occur at the oocyte stage, with poorly-performing oocytes potentially re-adsorbed. By contrast, MR-assisted IVF creates combinations of mito-nuclear alleles that are potentially novel (i.e. never before placed together), and not previously screened by natural selection (or previously screened and selected against). This lack of prior screening means that the sample of oocytes and embryos created under MR will contain individuals that may be inherently more likely to exhibit incompatibilities between the mitochondrial and nuclear genomes^8^. The mito-nuclear allelic combinations carried by the offspring will be under selection across life-stages, from before fertilization through to the sexually mature adult (with this selection manifested as differential patterns of survival or fertility among offspring carrying different mito-nuclear allelic combinations). Reduced fertility, especially of males, as a result of epistatic interactions during hybridization between alleles at different loci, including those spanning different genomes, is expected theoretically^11^ and supported empirically, including for mito-nuclear complexes in *Drosophila melanogaster* ^12,13^.

## 2. Genetic diversity in humans

The human population is generally thought to show lower mean levels of genetic divergence at nuclear loci than other species^e.g. 14^. While the probability of MR resulting in mito-nuclear incompatibilities would presumably be low if there was complete genetic admixture within the nuclear genome, it is clear that genetic population stratification does exist^15–17^. This stratification has its origins in historical and demographic patterns of selection and migration^18^, and positive assortative mating between individuals of similar phenotypes may contribute to its maintenance^19,20^.

However, the level of divergence across mtDNA sequences is also relevant when it comes to the question of whether or not MR may result in mismatched mito-nuclear genotypes. In humans, the percentage divergence in mtDNA between major human haplogroups is around 0.5% (Fig 1, Table S1), essentially equivalent to the divergence exhibited across mtDNA haplotypes within the fruit fly, *D. melanogaster* (0.4%; Fig 1, Table S2), which exhibit clear signatures of mito-nuclear incompatibilities, particularly in males^12,13^. Haplogroup matching, proposed as a way of circumventing this issue^21,22^, mig ht not always be successful in preventing mito-nuclear incompatibilities. By definition, when probing variation within human macro-haplogroups, divergence across mtDNA haplotypes will persist (∼0.1%, see Table S3 for estimates within haplogroup H, the most common European macro-haplogroup, or 0.2% if the non-coding region is included in the analysis [Fig 1]; similar patterns are found within H1 [Table S4, Fig 1]). The mechanisms of the incompatibilities are largely unknown and the identity of the causative interacting loci is undetermined^23^. However, it seems that several loci of small effect are involved in *Drosophila*^24^. While this suggests that less distantly-related genomes may result in smaller incompatibility effects^24^, it will be difficult to make predictions about the likelihood of incompatibilities based on the specific alleles that delineate haplotypes, given that it has been previously shown that single nucleotide differences in the mtDNA can cause male sterility when interacting with particular nuclear genotypes^13,25^. Further research into the degree of mismatch manifested with increasing mitochondrial genetic divergence between putative donor and patients should be a priority.

**Figure 1:**
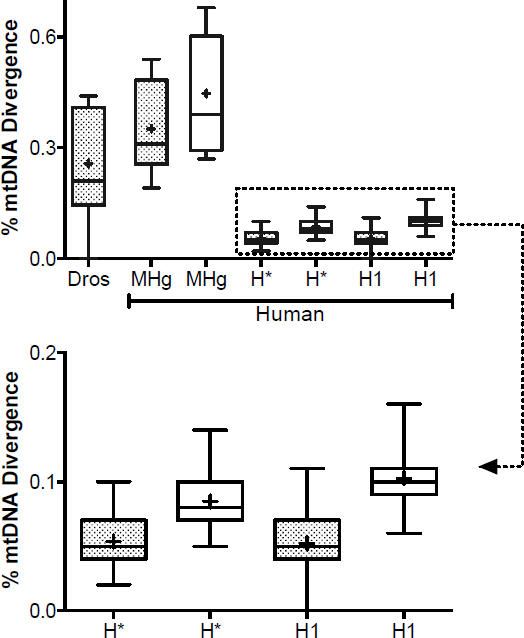
Boxplots depicting variation in mtDNA divergence (%) across naturally-occurring human mtDNA sequences, with comparison to mtDNA divergence across global fruit fly (*Drosophila melanogaster*) populations. The *Drosophila* (Dros) plot is based on protein coding regions of 13 *Drosophila melanogaster* populations that were previously used in published studies showing effects of mitochondrial replacement^12,13,24^. Human data are first presented using sequence polymorphisms found only in the protein coding region (denoted by hashed boxes; to enable direct comparison to the Drosophila plots, in which non-protein coding sequences were unavailable), and secondly using the full sequence data (protein and non-coding regions; denoted by open boxes). Human data are presented at three scales; first at the scale of human mitochondrial macro haplogroups M, N, R, L0, and L3 (MHg), second at the scale of human mitochondrial haplogroup H (sub-clades H1 to H10 [H*]) – the most common haplogroup among Europeans, and third at the scale of haplotype, specifically 20 mitochondrial haplotypes sampled from haplogroup H1 (H1). Box plots show median values (line within box), 2^nd^, and 3^rd^ quartile (box outline), maximum data range (whiskers), and mean (+). At each scale, plots are generated using pairwise divergence estimates for all combinations of mtDNA sequence.

## 3. Proof-of-principle studies do not allow epidemiological predictions of incompatibilities

Several studies demonstrated the technical feasibility of surgical MR^1–3,26^, using macaques, human cell lines and mice. The number of mitochondrial × nuclear genotype combinations covered by all these studies together appears to be 15, or less. In addition, with the exception of two studies^3,27^, no maternal replicates were used per mito-nuclear combination, preventing an examination of whether any effect is due to a particular maternal effect associated with the study subject or inherent to a particular mitochondrial × nuclear genotype combination. In other words, these studies^1–3,26,27^ were not designed to test for mito-nuclear incompatibilities, and cannot be used to predict the population-wide likelihood of mito-nuclear incompatibilities manifesting post-MR. Doing so will likely lead to a high rate of Type II errors – the failure to detect effects that are present.

There are also additional issues with some of these studies. For example, Tachibana et al.^3^ used 98 human oocytes (from 7 donors) for MR, and concluded, there was no difference in zygote survival to normal IVF controls. However, this conclusion might warrant reappraisal. In their study, the authors derived six embryonic stem cells (ESCs), from 19 blastocysts, from a starting stock of 64 oocytes that underwent MR treatment (ESC success rate: 6/64 = 9%, blastocyst rate: 19/64 = 30%). This compares to nine ESCs, from 16 blastocysts, from a starting stock of 33 oocytes in the control group (ESC rate: 9/33 = 27%, blastocyst rate: 16/33 = 48%). The differences in ESC isolation rate between treatment and control groups are in fact statistically significant (ESC: Fisher’s exact test, 1df, two-tailed, p = 0.035; Blastocysts: p = 0.078). This suggests further evaluation of developmental success post-MR should be a priority.

Paull et al.^26^ obtained 7 blastocysts out of 18 MR oocytes. The cell lines derived from these blastocysts showed lower activity in all four respiratory chain enzyme complexes than control cells. While differences were not statistically significant, they represent reductions of between 2 to 19 % (average across four enzymes: 11%) compared to parthogenetically-induced controls, and given the low sample sizes involved, again suggest that further scrutiny into possible effects of MR is warranted.

Finally, Craven et al^2^ report that development to blastocyst stage was approximately 50% lower for zygotes receiving MR treatment (18/80 = 22.5%) than for controls; a difference that is likely to be statistically significant, although the controls were unmanipulated and therefore do not represent a true control for the manipulation.

In the light of these three examples, it is noteworthy, that MR affected development and respiration in many other studies on non-primate vertebrates and invertebrates^8^.

## Conclusions

MR-assisted IVF could place novel allelic combinations of interacting mtDNA and nuclear genes alongside each other in the offspring, and these combinations might not have been previously screened by selection (or in the worst case may have already been removed from the population by selection). Therefore, mito-nuclear allelic combinations created following MR (which are characterized by 0% co-transmission from parents to offspring) are theoretically not equivalent to those found in individuals produced under sexual reproduction. This insight is of fundamental importance, but apparently underappreciated in the literature pertaining to MR. Given that mito-nuclear allelic combinations contribute to encoding life’s critical function of energy conversion, natural selection must be assumed to be particularly intense on these combinations. We suggest that it is a real possibility that novel combinations created under MR could result in mito-nuclear mismatches. This possibility has also been predicted by evolutionary theory^28^ and experimentally supported in several taxa^8,29^, including several with comparable levels of mitochondrial genetic diversity to the human population^12,13^.

Lack of evidence from small-scale proof-of-principle experiments for MR effects should not be used to conclude mito-nuclear incompatibilities are unlikely to manifest post-MR, because these experiments cover few mito-nuclear combinations and their statistical inferences, in some cases, appear open to question. In fact, there is actually an extensive, but largely overlooked, body of experimental evidence that indicates mito-nuclear interactions are important in determining health outcomes in humans^30–42^, as well evidence for mito-nuclear incompatibilities following the similar procedure of somatic cell nuclear transfer in cattle^43,44^. Furthermore, the only previous attempt of using pronuclear transfer in humans was not successful^45^. Future work should, therefore, address to what extent the risk of mismatching can be reduced by matching the donor and maternal mitochondrial haplotypes, since genetic variation across many interacting loci are likely to be involved^24^, and given the genetic variation between and within human mtDNA haplogroups that we have outlined here. As a suggested design, two oocytes should be used for every donor; each enucleated. One of these is assigned to a control, and re-populated with the donor’s own nuclear genetic material, and the other to the MR treatment. By then comparing the success of MR-treated to control eggs, and provided sufficient replication across donors, this design would provide an explicit test for mito-nuclear incompatibilities post-MR.

## Acknowledgements

Funding was provided by Royal Society University Research Fellowship and European Research Council (to EHM), the VolkswagenFoundation and the Zukunftskonzept at TU Dresden funded by the Exzellenzinitiative of the Deutsche Forschungsgemeinschaft (to KR), and the Australian Research Council (to DKD).

